# Cellulose biosynthesis inhibition reduces cell cycle activity in a nitrate reductase- and cytokinin-dependent manner

**DOI:** 10.1101/286161

**Authors:** Nora Gigli-Bisceglia, Timo Engelsdorf, Miroslav Strnad, Lauri Vaahtera, Amel Jamoune, Leila Alipanah, Ondřej Novák, Jan Hejatko, Thorsten Hamann

## Abstract

- During growth, development and defense, cell wall integrity needs to be coordinated with cell cycle activity. In *Saccharomyces cerevisiae,* coordination is mediated by the cell wall integrity maintenance mechanism. In plants, little is known how coordination is achieved.
- Here we investigated coordination between plant cell wall and cell cycle activity in *Arabidopsis thaliana* seedlings by studying the impact of cell wall damage (CWD, caused by cellulose biosynthesis inhibition) on cell cycle gene expression, growth, phytohormone (jasmonic acid, salicylic acid, cytokinins) and lignin accumulation.
- We found root growth and cell cycle gene expression are reduced by CWD in an osmo-sensitive manner. *trans*-zeatin application suppressed the CWD effect on gene expression. Quantification of cytokinins revealed CWD-induced, osmo-sensitive changes in several cytokinins. Expression of *CYTOKININ OXIDASE2/DEHYDROGENASE* (*CKX2*) and *CKX3*, encoding cytokinin-degrading enzymes, was elevated in CWD-exposed seedlings. Genetic studies implicated *NITRATE REDUCTASE1/2* (*NIA1/2*) in the response to CWD. In *nia1/2* seedlings CWD induced neither expression of *CKX2/3* and cell cycle genes nor accumulation of jasmonic acid, salicylic acid and lignin.
- This suggests that CWD causes increased *CKX2*/*3* expression through a *NIA1/2*-mediated process. Increased *CKX* expression seems to cause changes in cytokinin levels, leading to reduced cell cycle gene expression.

## Introduction

The chemically complex cell walls surrounding all plant cells are a distinctive feature of plants. Walls provide rigidity during defense and are plastic during growth, cell morphogenesis and division. This dual function is achieved by actively modulating wall metabolism to modify the carbohydrate-based complex wall structure consisting mainly of cellulose, hemicelluloses, pectins, lignin and different cell wall proteins (Höfte & Voxeur, 2017). Despite the obvious necessity for coordination between cell wall metabolism and cell division, knowledge regarding the underlying mechanism is limited. In *Saccharomyces cerevisiae,* the cell wall integrity (CWI) maintenance mechanism is intricately involved in this coordination (Levin, 2011). A dedicated checkpoint monitoring yeast CWI, regulates cell cycle progression depending on the state of the wall (Levin, 2011; Kono *et al*., 2016).

In plants, CWI is impaired during growth and following exposure to abiotic and biotic stress (Wolf, 2017; Bacete *et al*., 2018; Feng *et al*., 2018). CWI-dependent responses include modifications of cellular metabolism (such as osmo-sensitive changes in carbohydrate metabolism and phytohormone production), growth inhibition, modification of pectic polysaccharides and deposition of lignin and callose (Ellis & Turner, 2001; Cano-Delgado *et al*., 2003; Hematy *et al*., 2007; Hamann *et al*., 2009; Tsang *et al*., 2011; Wormit *et al*., 2012; Zhao & Dixon, 2014; Bacete *et al*., 2018). Currently, knowledge regarding the plant CWI maintenance mechanism is still very limited. Osmotic support (mannitol, poly-ethylene glycol or sorbitol) can suppress most responses induced by cell wall damage (CWD) in a concentration dependent manner both in yeast and Arabidopsis seedlings, suggesting that similarities might exist between the two organisms (Hamann *et al*., 2009; Levin, 2011; Hamann, 2015). This is further supported by the observation that ARABIDOPSIS HISTIDINE KINASE1 (AHK1) partially complements yeast strains deficient in osmosensing and is required for osmo-perception and abiotic stress responses in plants (Tran *et al*., 2007). In addition, MID1-COMPLEMENTING ACTIVITY1 (MCA1) from Arabidopsis partially complements a yeast strain deficient in the plasmamembrane-localized Ca^2+^-channel protein complex MID1-CCH1, involved in the yeast CWI maintenance mechanism (Paidhungat & Garrett, 1997; Nakagawa *et al*., 2007; Levin, 2011). In Arabidopsis, MCA1 mediates Ca^2+^-based signaling processes, CWI maintenance and both hypo-osmotic stress and mechano-perception (Yamanaka *et al*., 2010; Denness *et al*., 2011; Furuichi *et al*., 2016). Ca^2+^ signaling inhibitors prevent CWD-induced phytohormone (jasmonic acid, JA; salicylic acid, SA) accumulation and lignin production in Arabidopsis seedlings, implicating Ca^2+^-based signaling processes (Denness *et al*., 2011). Several *Catharanthus roseus* receptor like kinases (CrRLK), have been implicated in CWI maintenance (Engelsdorf & Hamann, 2014). THESEUS1 (THE1) is required during CWI maintenance and resistance to *Fusarium oxysporum* f. sp. *conglutinans* infection (Hematy *et al*., 2007; Van der Does *et al*., 2017), while FERONIA (FER) is required for CWI maintenance during salt stress (Feng *et al*., 2018). Homologs for MCA1 in *Zea mays* (NOD), *Oryza sativa* (OsMCA1)) and THESEUS1 in *Marchantia polymorpha* (MpTHE) have been identified, suggesting conservation of the CWI maintenance mechanism (Kurusu *et al*., 2012; Honkanen *et al*., 2016; Rosa *et al*., 2017).

In many cell wall biosynthetic mutants growth defects are observed, which are possibly caused by hyperactive defense signaling triggered by cell wall defects (Bacete *et al*., 2018). Dwarf phenotypes have also been associated with loss of anisotropic cell expansion due to mutations in *CELLULOSE SYNTHASEs* (*CESAs*) encoding proteins required for cellulose biosynthesis during cell wall formation and cell division (McFarlane *et al*., 2014; Chen *et al*., 2018). Cellulose biosynthesis inhibitors have become routinely used tools, since they allow CWD generation in a tightly controlled manner (temporally and spatially), enabling targeted studies of early and late stress responses and biosynthetic effects (Manfield *et al*., 2004; Wormit *et al*., 2012; Tateno *et al*., 2015) One of the inhibitors, isoxaben (ISX), inhibits specifically cellulose production during primary cell wall formation and affects the subcellular localization of the cellulose synthase complexes within minutes of application (Heim *et al*., 1990; Paredez *et al*., 2006; Gutierrez *et al*., 2009). ISX is frequently because of the availability of resistance causing mutations, like *isoxaben-resistant1* (*ixr1-1*), which represent efficient controls (Scheible *et al*., 2001). The *ixr1-1* mutation resides in *CELLULOSE SYNTHASE A3* (*CESA3*), which encodes a subunit of the rosette complex active during primary cell wall formation. ISX-triggered cellulose reduction, similar to cellulose deficient mutants, leads to changes in gene expression, loss of anisotropic cell expansion, ectopic lignification, JA and SA production, and ultimately cell death (Ellis *et al*., 2002; Cano-Delgado *et al*., 2003; Hamann *et al*., 2009). ISX-induced responses, including accumulation of phytohormones, generation of reactive oxygen species (ROS) and lignin, can be inhibited with diphenylene iodonium treatments (Denness *et al*., 2011). Diphenylene iodonium is an inhibitor of flavin-containing enzymes implicated in ROS-production, whose specificity is concentration-dependent but its precise mode of action in plants is unknown (Kärkönen & Kuchitsu, 2015). Another cellulose biosynthesis inhibitor (C17) was identified through its ability to inhibit cell division, showing that cellulose biosynthesis inhibition affects cell division (Hu *et al*., 2016).

Plant cell walls and the cell cycle undergo major changes in response to drought and osmotic stress (Skirycz *et al*., 2011; Tenhaken, 2014). While the response to osmotic stress is mediated by an ethylene-dependent mechanism, cytokinins are involved in drought stress adaptation both as positive and negative regulators (Nishiyama *et al*., 2011; Li *et al*., 2016; Nguyen *et al*., 2016). Interestingly, cytokinin application to plants and cell cultures induces expression changes in genes encoding cell wall modifying enzymes (expansins, laccases and pectin-modifying enzymes), suggesting that cytokinins can modify plant cell wall metabolism (Brenner *et al*., 2012). While the precise mode of action of cytokinins remains to be determined, Dirigent-like proteins could form a regulatory element (Paniagua *et al*., 2017). Their expression is regulated both by cytokinins and different biotic stresses and they mediate stereo-selectivity during lignan biosynthesis. Cytokinins regulate also plant cell cycle progression (Schaller *et al*., 2014). They modulate transition from G2- to M-phase by regulating CDKA/B and CYCB1 activity and from G1- to S-phase by controlling *CYCD3;1* expression (Scofield *et al*., 2013). Cytokinin amounts in turn are regulated in different ways, including through changes in nitric oxide (NO) levels (Zürcher & Müller, 2016). NO can interact directly with *trans*-zeatin (*t*Z) type cytokinins creating nitrated cytokinin species and lower both NO and *t*Z-levels or inhibit phosphorylation of cytokinin signaling components (Feng *et al*., 2013; Liu *et al*., 2013). Cytokinins also regulate NO levels, which in turn control *CYCD3;1* induction, suggesting the existence of a regulatory loop involving both (Tun *et al*., 2008; Shen *et al*., 2013). Mutations in *NITRATE REDUCTASE 1* and *NITRATE REDUCTASE 2* (*NIA1/2*), impairing nitrate assimilation and NO production, form useful genetic tools to study NO-based signaling (Tun *et al*., 2008; Chamizo-Ampudia *et al*., 2017).

Here we combined phenotypic and genetic characterizations, with gene expression analysis and cytokinin measurements to investigate if and how cell cycle activity in *Arabidopsis thaliana* seedlings is coordinated with CWI. The results suggest that a *NITRATE REDUCTASE 1/2* (*NIA1/2)*-dependent, turgor-sensitive process, controlling expression of cytokinin degrading enzymes is responsible for changes in certain cytokinin levels in response to CWD. The changes in cytokinin levels seem to attenuate cell cycle activity by reducing *CYCD3;1* expression.

## Material and Methods

### Plant growth and treatment

*Arabidopsis thaliana* genotypes used are listed in Supplemental Table S1. Seedlings were sterilized and grown in liquid culture with minor modifications (Hamann *et al*., 2009). 30 mg of seeds were sterilized by sequential incubation with 70 % ethanol and 50 % bleach on a rotating mixer for 10 min each and washed 3 times with sterile water. All experiments were performed in 6 days old seedlings treated with 600 nM isoxaben (ISX; in DMSO), 300 mM sorbitol (s300) or a combination of both, in freshly prepared medium. Zeatin (1 mM stock solution) (Sigma-Aldrich) was dissolved in water. Seedlings were grown in long-day conditions (16 h light, 22°C / 8 h dark, 18°C) at 150 µmol m^-2^ s^-1^ photon flux density on a IKA KS501 flask shaker at a constant speed of 130 rotations per minute.

### Glucuronidase (GUS) staining

GUS staining was performed using staining buffer containing 50 mm sodium phosphate (pH 7.0), 0.5 mm potassium ferrocyanide, 10 mm EDTA, 0.1% (v/v) Triton X-100, 2% (v/v) dimethyl sulfoxide, and 1 mm 5-bromo-4-chloro-3-indolyl-d-glucuronide at 37°C for 24 h (Jefferson *et al*., 1987; Savatin *et al*., 2014).

### Analysis of root growth phenotypes

Root length measurements were performed right before (0 h) and 24 h after treatments (DMSO/mock, ISX, s300 or ISXs300). Absolute RAM (root apical meristem) length was determined in primary roots treated as before for 3, 6 and 9 h. Root tips were stained for 5 minutes with propidium iodide (1 mg/ml), followed by two washing steps with MilliQ water and imaging using a LSM Zeiss 710. RAM length (defined as the distance between the quiescent center (QC) and the first elongating cell in the root cortical cell layer) and RAM size (expressed as number of cortical cells in the RAM) were measured using FIJI (https://fiji.sc).

### Cytokinin measurements

Quantification of cytokinin metabolites was performed according to a method described previously (Svačinová *et al*., 2012) and slightly modified as described before (Antoniadi *et al*., 2015). Samples were homogenized and extracted in 1 ml of modified Bieleski buffer (60% MeOH, 10%HCOOH and 30%H_2_O) together with a cocktail of stable isotope-labeled internal standards (0.25 pmol of Cytokinin bases, ribosides, *N*-glucosides and 0.5 pmol of Cytokinin *O*-glucosides per sample). Extracts were purified using a Oasis MCX column (30 mg/1 ml, Waters) conditioned with 1 ml each of 100% MeOH and H_2_O, equilibrated sequentially with 1 ml of 50% (v/v) nitric acid, 1 ml of H_2_O, and 1 ml of 1 M HCOOH. After sample application onto an MCX column, un-retained compounds were removed by a wash step using 1 ml of 1 M HCOOH and 1 ml 100% MeOH, pre-concentrated analytes were eluted by two-step elution using 1 ml of 0.35M NH_4_OH aqueous solution and 2 ml of 0.35 M NH_4_OH in 60% (v/v) MeOH solution. Eluates were evaporated to dryness *in vacuo* and stored at -20°C. Cytokinin levels were determined using ultra-high performance liquid chromatography-electrospray tandem mass spectrometry (UHPLC-MS/MS) (Rittenberg D., 1940).

### Salicylic acid and jasmonic acid quantification

JA and SA were analyzed as described by (Forcat et al. 2008) with minor modifications. Briefly, 6-7 mg freeze-dried seedlings were ground in a Qiagen Tissue Lyser II and extracted with 400 µl extraction buffer (10 % methanol, 1 % acetic acid) containing internal standards (10 ng Jasmonic-d_5_ Acid, 28 ng Salicylic-d_4_ Acid; CDN Isotopes, Pointe-Claire, Canada). Samples were incubated on ice for 30 min and cell debris pelleted by centrifugation. Pellets were re-extracted with 400 µl extraction buffer without internal standards, supernatants combined and used for LC-MS/MS analysis. Mass transitions were: JA 209 > 59, D_5_-JA 214 > 62, SA 137 > 93, D_4_-SA 141 > 97.

### qRT-PCR

Total RNA was isolated using a Spectrum Plant Total RNA Kit (Sigma-Aldrich). 2 μg of total RNA were treated with RQ1 RNase-Free DNase (Promega) and cDNA synthesis was performed with the ImProm-II Reverse Transcription System (Promega). qRT-PCR was performed using a LightCycler 480 SYBR Green I Master (Roche) and primers (Supplemental Table S2) diluted according to manufacturer specifications. *ACT2* was used as reference gene.

### Transcriptomic analysis

Total RNA was extracted using a Spectrum Plant Total RNA Kit (Sigma-Aldrich) from three biological replicates for Col-0 and two for *nia1 2* for both treatments. RNA integrity assessed using a Agilent RNA 6000 Pico Kit. Sequencing samples were prepared as described before (Ren *et al*., 2015). Single read sequencing was performed for 75 base pair read length on a NextSeq500 instrument (Illumina RTA v2.4.6). FASTQ files were generated using bcl2fastq2 Conversion Software v1.8.4. Each FASTQ file was subjected to quality control trough fastQC v11.1 before technical replicates were combined and an average of 13.1 million reads was produced for each library. Reads were aligned to the *A. thaliana* genome (Ensembl v82) with STAR v2.4.1 in two-pass mode. On average, 96.2% of the reads aligned to the genome. Reads that aligned uniquely to the genome were aggregated into gene counts with Feature Counts v1.4.6 using the genome annotations defined in Ensembl v82. Differentially expressed genes were extracted from Col-0 and *nia1 nia2* genotypes after ISX/mock treatments using DESeq2 differential expression analysis pipeline (Love *et al*., 2014). For both genotypes, genes showing differential expression in response to ISX treatment vs. mock with a 1% False Discovery Rate (FDR) were considered significantly differentially expressed (DE). In order to assess the variance within the samples, a principal component analysis (PCA) was performed. Regularized logarithm transformation from DESeq2 package was applied to read counts and PCA was performed for 2000 genes with highest variance using “prcomp” function. The intersection of the DE genes between the genotypes was visualized using a Venn diagram produced with Venny 2.1 (http://bioinfogp.cnb.csic.es/tools/venny/index.html). Analysis of GO categories was performed using AtCOECiS (Vandepoele *et al*., 2009). Transcriptomics data has been deposited with the gene expression omnibus under the following submission ID: GSE109613.

### Statistical Analysis

Statistical significance (Student’t test, ANOVA) was assessed using IBM SPSS Statistics v24.

## Results

### Osmotic treatments attenuate ISX-derived root growth inhibition

To analyse the regulatory mechanism coordinating CWI and cell cycle progression we determined first when and how cellulose biosynthesis inhibition affects *Arabidopsis thaliana* seedling root growth and root apical meristems (RAM) organization. We used an established model system where seedlings are grown under controlled conditions in liquid culture and cellulose production is inhibited by isoxaben (ISX) application (Hamann *et al*., 2009). Initially, we confirmed ISX-specificity by determining root lengths in mock- or ISX-treated Col-0 and ISX-resistant *ixr1-1* seedlings (Fig. 1a). Mock-treated (DMSO, solvent of ISX stock solution) Col-0 and *ixr1-1* seedlings exhibited similar root lengths both at start of treatment (T0) and after 24h. Root growth was nearly completely inhibited in Col-0 seedlings by ISX-treatment while no inhibition was detectable in *ixr1-1* seedlings, confirming ISX specificity. Osmoticum treatments (sorbitol, 300mM, s300) were included for the experiments since they have been shown to suppress ISX-induced responses in a concentration-dependent manner (Hamann *et al*., 2009).

**Figure 1.**
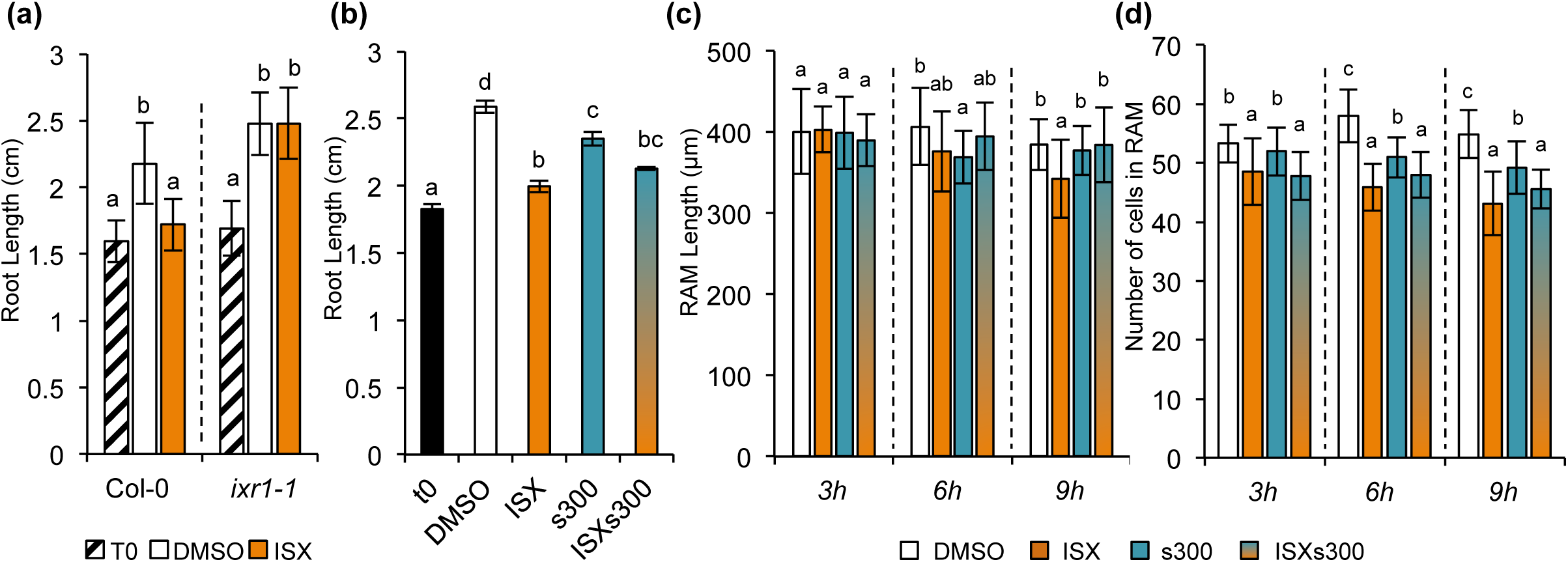
ISX-treatments induce sorbitol-sensitive root growth phenotypes. **a**) Root length was determined in Col-0 and *ixr1-1* seedlings (T0) and after 24 h of treatment with DMSO (mock) or Isoxaben (ISX). **b**) Root lengths in Col-0 seedlings at start of treatment (T0) or after 24h of treatment with DMSO, Isoxaben (ISX), Sorbitol (s300) or ISXs300. **c**) Seedlings were treated as described in b for 3, 6 and 9 h and RAM length was measured. **d**) RAM cell number was determined for seedlings treated as in b. 25 roots were analyzed per treatment, experiments were repeated three times and values represent averages of the three experiments. Error bars are based on standard deviation and different letters indicate statistically significant differences according to one-way ANOVA and Tukey’s HSD test (α = 0.05). In (a) ANOVA was used to analyze differences between the treatments within genotypes, in (b) to analyze differences within the treatments while in (c) and (d) to analyze the differences within single time points.

Seedlings were treated with DMSO (mock), ISX, s300 or a combination of ISX and sorbitol (ISXs300) and root lengths were measured as before (Fig. 1b). Root growth was reduced by s300 while ISX treatment caused a nearly complete stop. ISX/s300 treatment resulted in an intermediate (not additive) growth phenotype. To determine if and when treatment effects are detectable on the meristem level we measured both RAM length as well as cell number in seedling roots treated as before for 3, 6 or 9h. ISX-treatment reduced RAM length (Fig. 1c) significantly after 9 h and cell number after 6h (Fig. 1d). While s300 treatments had no consistent effects on RAM length, RAM cell number was consistently reduced. ISXs300 treatments resulted in RAM lengths similar to mock while RAM cell numbers were similar to ISX. These results suggest that ISX treatments affect both cell division and elongation. The osmotic support (s300) reduces the inhibitory effect on cell elongation but not on cell number indicating differential control of cell extension and division.

### Osmotic co-treatments attenuate ISX-triggered inhibition of cell cycle gene expression

To investigate if the treatments with ISX and osmoticum affect cell cycle gene expression we performed time course experiments as before with seedlings expressing a *pCYCB1;1::CYCB1;1 D-box-GUS* reporter (Colón-Carmona *et al*., 1999). GUS staining analysis suggested that *pCYCB1;1::CYCB1;1 D-box-GUS* transcription is strongly reduced in RAMs after 9 h of ISX-treatment compared to mock (Fig. 2a). s300 had milder effects on reporter expression (Fig. 2a). Reporter activity in RAMs exposed to ISXs300 seemed more similar to s300- than to ISX-treated ones (Fig. 2a). This effect is not restricted to the RAM since the reporter exhibited similar expression changes also in shoot apical meristems (SAMs) after 9 h (Fig. 2a). Because this analysis is not quantitative, qRT-PCR-based expression analysis was performed for three genes regulating different transition steps in cell cycle progression. The first one *CYCB1;1* is regulating G2- to M-transition, while the second one *CDKB2;1* is active in the late S- to M-phase (Okushima *et al*., 2014; Schaller *et al*., 2014) and the third one *CYCD3;1* is required during G1/S transition (Scofield *et al*., 2013). The genes examined exhibited dynamic transcript level changes with pronounced differences becoming apparent after 9 h (Fig. 2b). *CYCB1;1*, *CDKB2;1* and *CYCD3;1* transcript levels increased over time in mock-treated seedlings with slightly different dynamics (Fig. 2b). Transcript levels changed similarly in s300-treated seedlings (Fig. 2b).

**Figure 2.**
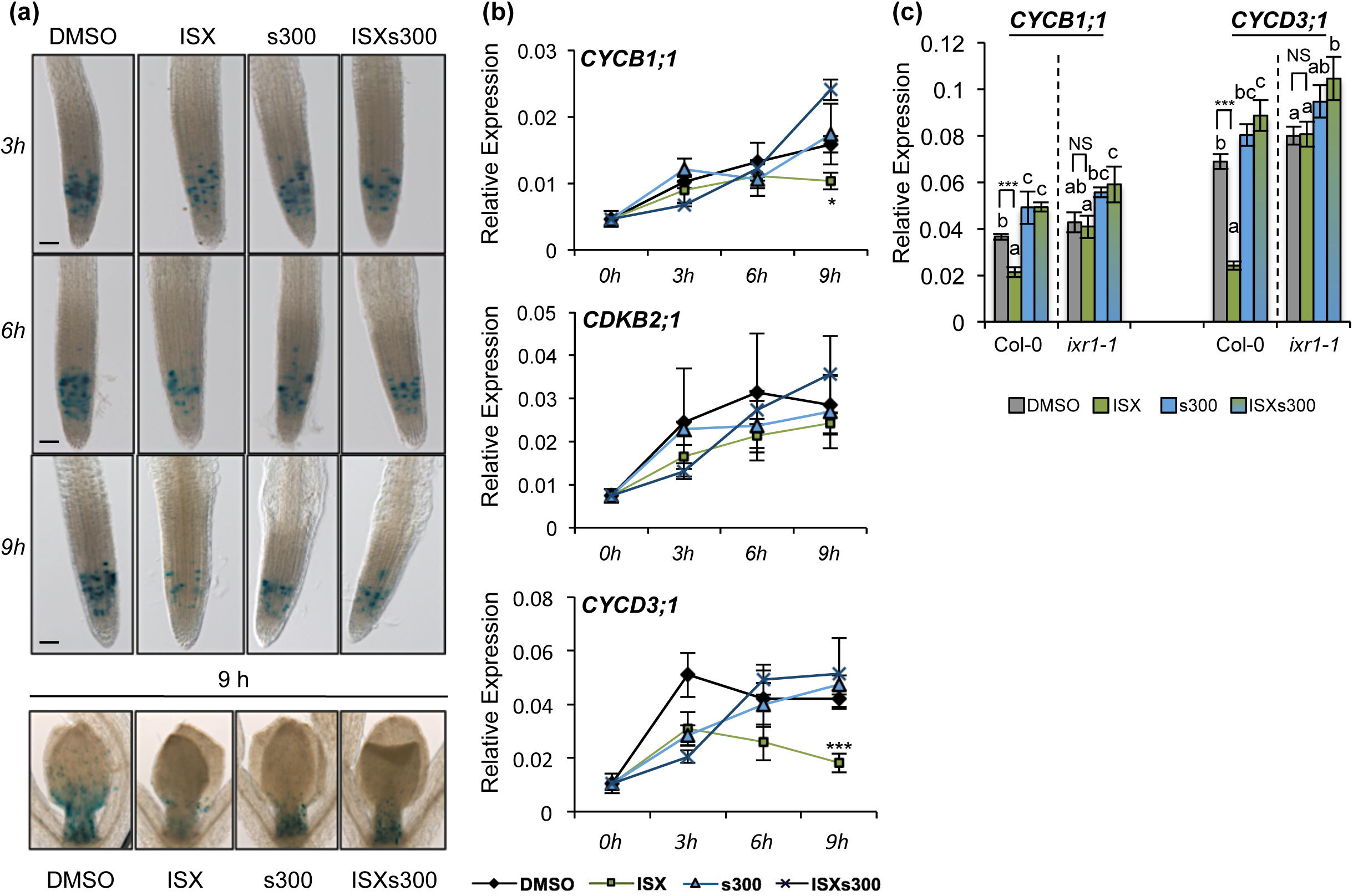
ISX-induced reduction of CYCB1;1, CYCD3;1 and CDKB2;1 transcript levels is attenuated by sorbitol co-treatments. **a**) *pCYCB1;1::CYCB1;1:GUS* seedlings were treated with DMSO (mock), Isoxaben (ISX), Sorbitol (s300) or a combination of both (ISXs300), collected after 3, 6 or 9 h and stained for GUS activity. **b**) Col-0 seedlings were grown and treated as in (a). Relative expression of *CYCB1;1*, *CDKB2;1* and *CYCD3;1* was quantified with qRT-PCR. **c**) Col-0 and *ixr1-1* seedlings were treated for 9 h as in (a) and gene expression analysed as in (b). Different letters denote significant differences according to one-way ANOVA and Tukey’s HSD test (α = 0.05) between different treatments of each genotype. In (b) and (c), values are means and error bars are based on standard deviation (n=3). Pairwise comparisons show select statistical differences for *CYCB1;1* or *CYCD3;1* transcript levels (in Col-0 and *ixr1-1)* in (c). Asterisks indicate differences according to Student’s t-test. * P < 0.05, *** P <0.001, ns = not significant.

Transcript levels increased early on also in ISX-treated seedlings. However, after three hours *CYCD3;1* transcript level started to decrease while *CYCB1;1* plateaued and *CDKB2;1* kept increasing slightly (Fig. 2b). *CYCD3;1* and *CYCB1;1* transcript levels were significantly lower in ISX- than mock-treated seedlings after 9 h, while *CDKB2;1* transcript levels did not differ between both treatments. In ISXs300-treated seedlings transcript levels of the three genes examined were not significantly different from mock controls after 9 h. To investigate whether the transcriptional effects observed depend on ISX-induced cellulose biosynthesis inhibition, Col-0 and *ixr1-1* seedlings were treated in the same manner. *CYCB1;1* and *CYCD3;1* transcript levels were investigated after 9 h, since those two genes exhibited the most pronounced changes. ISX-treated *ixr1-1* seedlings exhibited no transcript changes compared to mock while the changes caused by s300 were similar to Col-0 (Fig. 2c). These results established that the ISX-effects observed are caused by cellulose biosynthesis inhibition. The experiments performed so far used whole seedlings for RNA isolation. To investigate whether the transcriptional effects are tissue-specific (i.e. confined to SAMs or RAMs), experiments were repeated as before but either whole seedlings, shoots or roots only were used for RNA isolation and transcript levels of *CYCB1;1* and *CYCD3;1* were determined (Sup. Fig. 1). Transcript levels of both genes changed in a similar manner as observed on the whole seedling level suggesting that the effects observed are not restricted to a particular tissue.

Transcript levels for all genes examined increase over time in mock-treated seedlings possibly because the growth medium is replaced with fresh one at the start of the experiments (Riou-Khamlichi *et al*., 2000). Transcript level differences become pronounced after 9h. In ISX-treated seedlings *CDKB2;1* transcript levels are not significantly different whereas *CYCB1;1* are slightly and *CYCD3;1* pronouncedly reduced compared to mock controls. ISX-effects on transcript levels of selected cell cycle-associated genes are apparently similarly sensitive to osmotic manipulation as reported before for other ISX-induced responses (Hamann *et al*., 2009; Wormit *et al*., 2012). In all experiments *CYCD3;1* transcript levels seem to be most sensitive to the manipulations performed.

### ISX-induced changes in transcript levels require nitrate reductase activity

THE1 has been shown to mediate ISX-induced CWD responses and CWI maintenance (Hematy *et al*., 2007; Hamann *et al*., 2009). Therefore we investigated if the THE1-based CWI maintenance mechanism is also active here by characterizing KOs for RLKs *(THE1*, *WALL ASSOCIATED KINASE 2* (*WAK2*), *MDIS1-INTERACTING RECEPTOR LIKE KINASE 2* (*MIK2*), *FEI2*); ion channels *MSCS-LIKE 2/3* (*MSL2/3*) and *MCA1* (Kohorn *et al*., 2006; Hematy *et al*., 2007; Xu *et al*., 2008; Haswell *et al*., 2008; Denness *et al*., 2011; Van der Does *et al*., 2017). Moreover, since JA signaling has been also implicated in CWI maintenance and regulation of *CYCB1;1* expression, we included a KO allele for a key JA biosynthetic enzyme (*ALLENE OXIDE SYNTHASE*, *AOS*) (Ellis & Turner, 2001; Świątek *et al*., 2004). To investigate a possible involvement of NO-based signaling processes we used seedlings with mutations in *NIA1* and *NIA2*. We used the same experimental approach as before and quantified transcript levels after 9h. *CYCB1;1* and *CYCD3;1* transcript level changes induced by the different treatments were similar to Col-0 seedlings in *aos*, *mik2, wak2, fei2, the1* and *mca1* seedlings (Sup. Fig. 2a-c). In mock-treated *msl2/3* seedlings *CYCB1;1* and *CYCD3;1* transcript levels were elevated, but relative changes in the responses were similar to Col-0 (Sup. Fig. 2d). Transcript levels in mock-treated *nia1/2* seedlings were similar to Col-0, but ISX, s300 and ISX/s300 treatments did not cause changes in transcript levels (Fig. 3a). To ensure that the effects observed are not due to indirect consequences on growth caused by changes in nitrogen metabolism in *nia1/2* seedlings, we assessed the impact of the *nia1/2* mutations on seedling root growth and ISX-induced root growth inhibition (Fig. 3b). While root length was slightly reduced in mock-treated *nia1/2* seedlings, ISX treatment inhibited root growth in Col-0 and *nia1/2* seedlings similarly, suggesting loss of *NIA1/2* activity does not impair growth per se and ISX-induced root growth inhibition.

**Figure 3.**
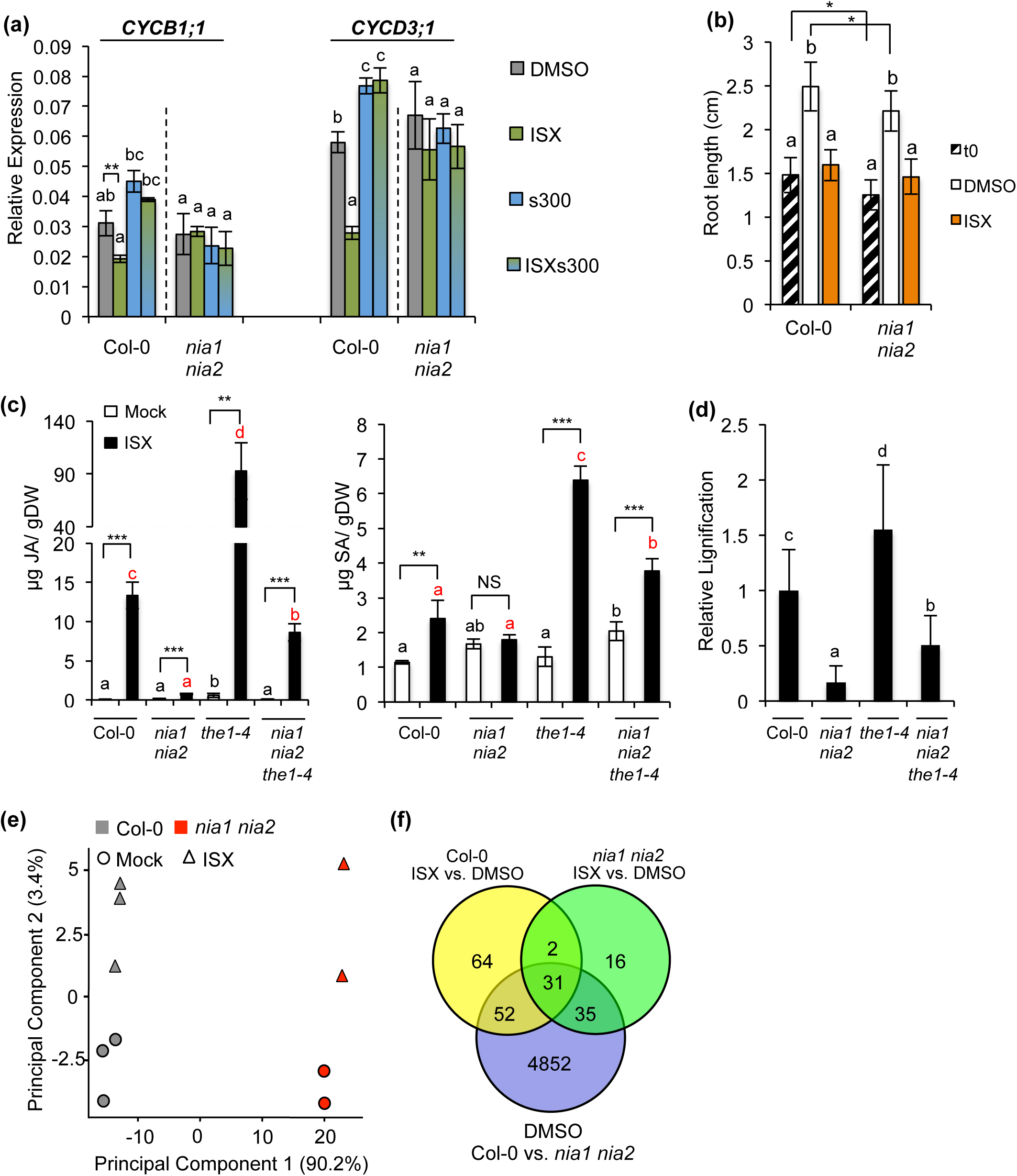
ISX-induced responses are reduced in nia1 nia2 seedlings. **a**) Col-0 and *nia1 nia2* seedlings were treated for 9 h with DMSO (mock), Isoxaben (ISX), Sorbitol (s300) or ISXs300. Relative expression of *CYCB1;1* and *CYCD3;1* was determined by qRT-PCR. **b**) Root lengths were determined in Col-0 and *nia1 nia2* seedlings (T0) and after 24 h of treatment with DMSO or ISX, Sorbitol (s300) or ISXs300. Values are means and error bars are based on standard deviation (n=25). **c**) Jasmonic acid (JA) and Salicylic acid (SA) accumulation was quantified in Col-0, *nia1 nia2, the1-4* and *nia1 nia2 the1-4* seedlings after 7 h of DMSO or ISX treatment. Values are means and error bars indicate standard deviation (n=3). Different letters denote significant differences. Pairwise comparisons were performed with Student’s t-test, * P < 0.05, ** P <0.01, *** P <0.001. **d**) Seedlings for indicated genotypes were stained with phloroglucinol HCl after 12 h and ectopic lignification was assessed by image analysis. Values are means and expressed relative to Col-0, with error bars indicating standard deviation (n= 20). **e**) Principal Component Analysis (PCA) of NGS data obtained from Col-0 and *nia1 nia2* seedlings after 1 h of ISX or mock treatment. **f**) Venn diagram showing differentially expressed genes significantly changing transcript abundance (P < 0.01) with data deriving from the same experiment as in (e). **a** - **d**) One-way ANOVA and Tukey’s HSD test (α = 0.05) were used to analyzedifferences between treatments of each genotype and different letters denote significant differences. In **b**) Comparisons were performed for the different genotypes upon mock (black letters) or ISX (red letters) treatment.

These results suggested that *NIA1/2* are involved in CWI maintenance and might be required for other *THE1*-mediated responses to ISX. To investigate this we introduced a *THE1* gain-of-function allele (*the1-4*) into the *nia1/2* background (Merz *et al*., 2017). We analyzed JA, SA and lignin accumulation in *nia1/2*, *the1-4* and *nia1/2 the1-4* seedlings after ISX-treatment. In ISX-treated *nia1/2* seedlings SA, JA and lignin production were dramatically reduced (Fig. 3c, d). While ISX-induced responses were enhanced in *the1-4* seedlings compared to Col-0, the same responses were reduced in *nia1/2 the1-4* seedlings compared to *the1-4* alone. This suggests that *NIA1/2*-dependent processes are required for ISX-induced JA, SA and lignin accumulation as well as full activation of THE1-mediated responses.

To understand the contributions of *NIA1/2* to ISX-induced responses, we performed a transcriptomics experiment with Col-0 and *nia1/2* seedlings mock- or ISX-treated for one hour. Initially a principal component analysis (PCA) was performed to assess the relative, global impact of genotypes and treatments on transcriptomes (Fig. 3e). The PCA showed that most of the differences result from the genotypes analyzed (PC1, 90.2%) whereas treatments had relatively minor effects (PC2, 3.4%). We then performed Gene Ontology (GO) analyses of the data. GO analysis of the results deriving from mock-treated Col-0 vs. *nia1/2* detected an over-representation of genes implicated in stress responses and primary metabolism (Sup. Table. S3). Analysis of Col-0 mock vs. Col-0 ISX and *nia1/2* mock vs. *nia1/2* ISX showed for both pronounced over-representation of defense response genes (Sup. Table S4, 5). The results of a more detailed data analysis are presented in a Venn diagram (Figure 3f). 4852 differentially expressed genes were detected between mock-treated Col-0 and *nia1/2*, underlining that *NIA1/2* contribute to fundamental processes. While 149 genes exhibited significant transcript level changes in ISX-treated Col-0 seedlings vs. mock, such changes were observed only for 84 genes in *nia1/2*. 33 genes exhibited significant differences in both ISX-treated Col-0 and *nia1/2* seedlings compared to mock controls, suggesting *NIA1/2* are not required for activation of these genes (Sup. Table S6). Amongst the 33 genes, several can be found, which encode components of Ca^2+^ signaling elements and transcription factors of ERF and WRKY families.

While the transcriptomic analysis detected fundamental differences between Col-0 and *nia1/2*, the growth assays and cell cycle gene expression analysis suggested limited effects of *nia1/2* on the processes examined here and indicate that *nia1/2* seedlings still respond to ISX. THE1-based CWI signaling and JA production are apparently not required for the ISX-induced reduction of *CYCB1;1* and *CYCD3;1* transcript levels. A *NIA1/2*-dependent process seems required for the ISX-induced changes in transcript levels, phytohormone / lignin accumulation and contributes to *THE1*-mediated processes.

### trans-zeatin application rescues ISX-triggered transcript level changes

Cytokinins regulate cell cycle progression (Schaller *et al*., 2014). To investigate if the ISX-effects on transcript levels are affected by cytokinin-based signaling processes, Col-0 seedlings were treated for 9h either with increasing concentrations of trans-zeatin (*t*Z) alone, ISX only or a combination of ISX and increasing *t*Z concentrations (Fig. 4a). All *t*Z treatments resulted in similarly increased *CYCB1;1* transcript levels in Col-0 seedlings. *CYCB1;1* transcript levels in Col-0 seedlings co-treated with ISX and *t*Z were inbetween transcript levels observed in ISX- or *t*Z-treated seedlings. *CYCD3;1* transcript levels were enhanced in a concentration dependent manner by *t*Z. Co-treatments with ISX and *t*Z led to *CYCD3;1* transcript levels similar to mock controls, showing a modifying effect of *t*Z on the ISX-derived reduction in *CYCD3;1* transcript levels. To test whether *NIA1/2* are required to mediate the effects of ISX and *t*Z treatments on transcript levels, we repeated the experiments with *nia1/2* seedlings (Fig. 4b). The transcript levels were similar in mock-treated *nia1/2* and Col-0 seedlings. *t*Z treatments induced elevated transcript levels at 100nM or 1 µM concentrations for both genes. Transcript levels for both genes were not reduced by ISX-treatments. In seedlings co-treated with ISX and 100nM or 1 µM *t*Z, transcript levels of both genes were similar to levels observed in seedlings treated with the corresponding *t*Z concentrations alone (no reduction detectable).

**Figure 4.**
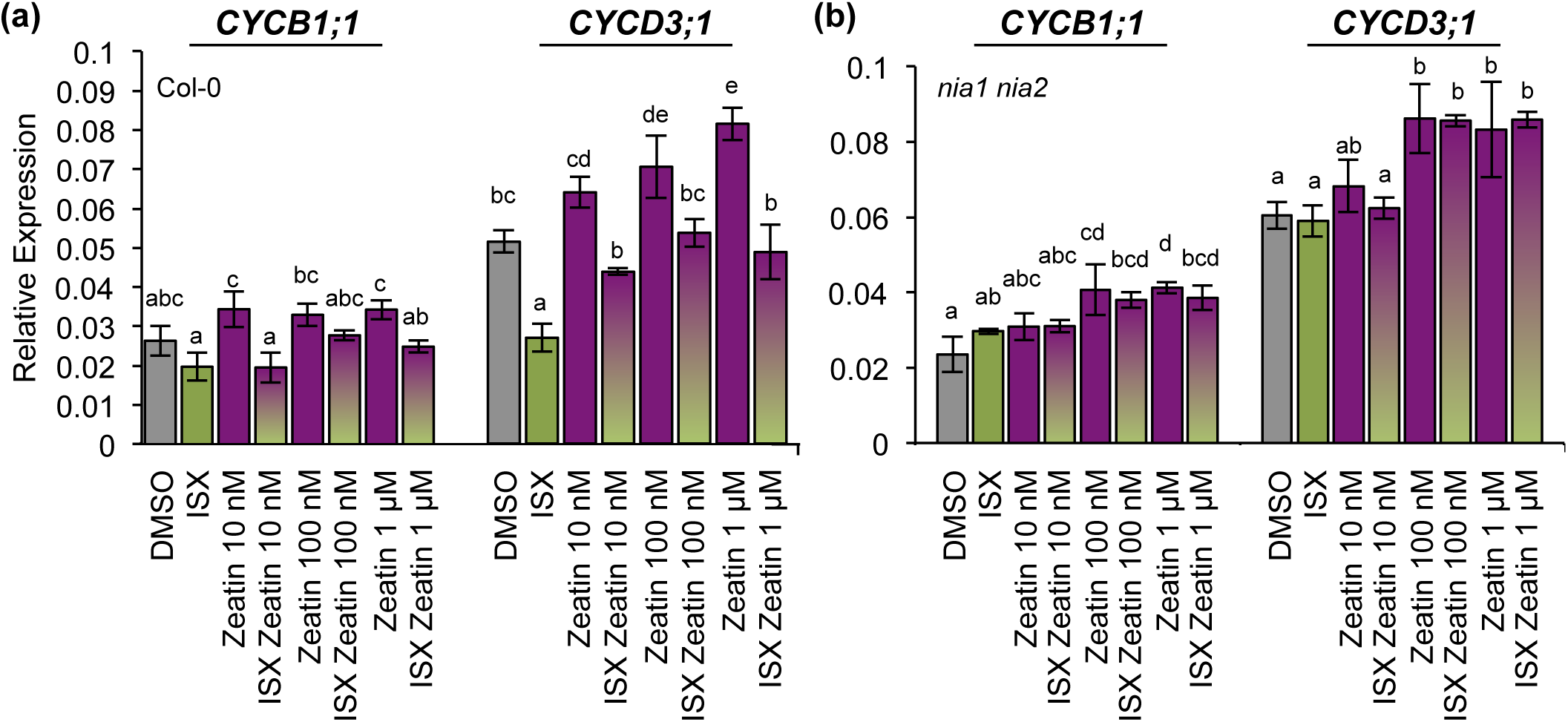
ISX-induced cell cycle transcript reduction is modulated by cytokinin application. Col-0 **a**) and *nia1 nia2* **b**) seedlings were treated with mock (DMSO), ISX, DMSO/zeatin and combinations of zeatin (10 nM, 100 nM and 1 µM) and ISX. Values are means and error bars are based on standard deviation (n=3). One-way ANOVA and Tukey’s HSD test (α = 0.05) were used to analyze the differences between treatments of each genotype and different letters denote significant differences.

To summarize, *t*Z treatments increase *CYCD3;1* transcript levels in a concentration dependent manner while effects on *CYCB1;1* transcripts are less pronounced in Col-0 seedlings. *t*Z application attenuates the ISX-effect on transcript levels in Col-0. In *nia1/2* seedlings, transcript levels are not reduced upon ISX treatment, while *t*Z treatments still resulted in elevated *CYCD3;1* and *CYCB1;1* transcript levels. This suggests the ISX-effects are dependent on *NIA1/2* while the *t*Z effect is independent of *NIA1/2*.

### Cytokinin measurements detect ISX-induced and osmo-sensitive changes in cytokinin levels

The data from the *t*Z experiments suggested that cytokinins modulate the ISX effects on *CYCD3;1* and *CYCB1;1* transcript levels. To determine if ISX treatments affect cytokinin levels, we measured total cytokinins and different isoprenoid cytokinins (incl. *t*Z, *cis*-zeatin (*c*Z), *N*^6^- (Δ^2^-isopentenyl) adenine (*i*P)) in Col-0 seedlings, mock (DMSO)-, ISX-, s300- or ISXs300- treated for up to 9 h (Fig. 5). Results for cytokinin bases, *cZ/t*Z and *i*P suggested type-specific changes after 3 h. Total CKs and *c*Z-/*t*Z-/*i*P-types started to show differences after 6 h, which became pronounced after 9 h. ISX-treated seedlings exhibited reduced levels of total Cytokinins, *t*Z- and *i*P-types while *c*Z-types were enhanced. Sorbitol treatments resulted in intermediate levels (between ISX and DMSO) with the exception of total *i*P-types where the levels were similar to those of mock-controls. Intriguingly, in ISXs300-treated seedlings levels of most cytokinin types were more similar to s300 than to ISX alone. To investigate if the changes observed might be tissue specific we also analyzed CK levels in aerial parts only after 9 h (Sup. Figure S3). The results showed a similar trend as in whole seedlings.

**Figure 5.**
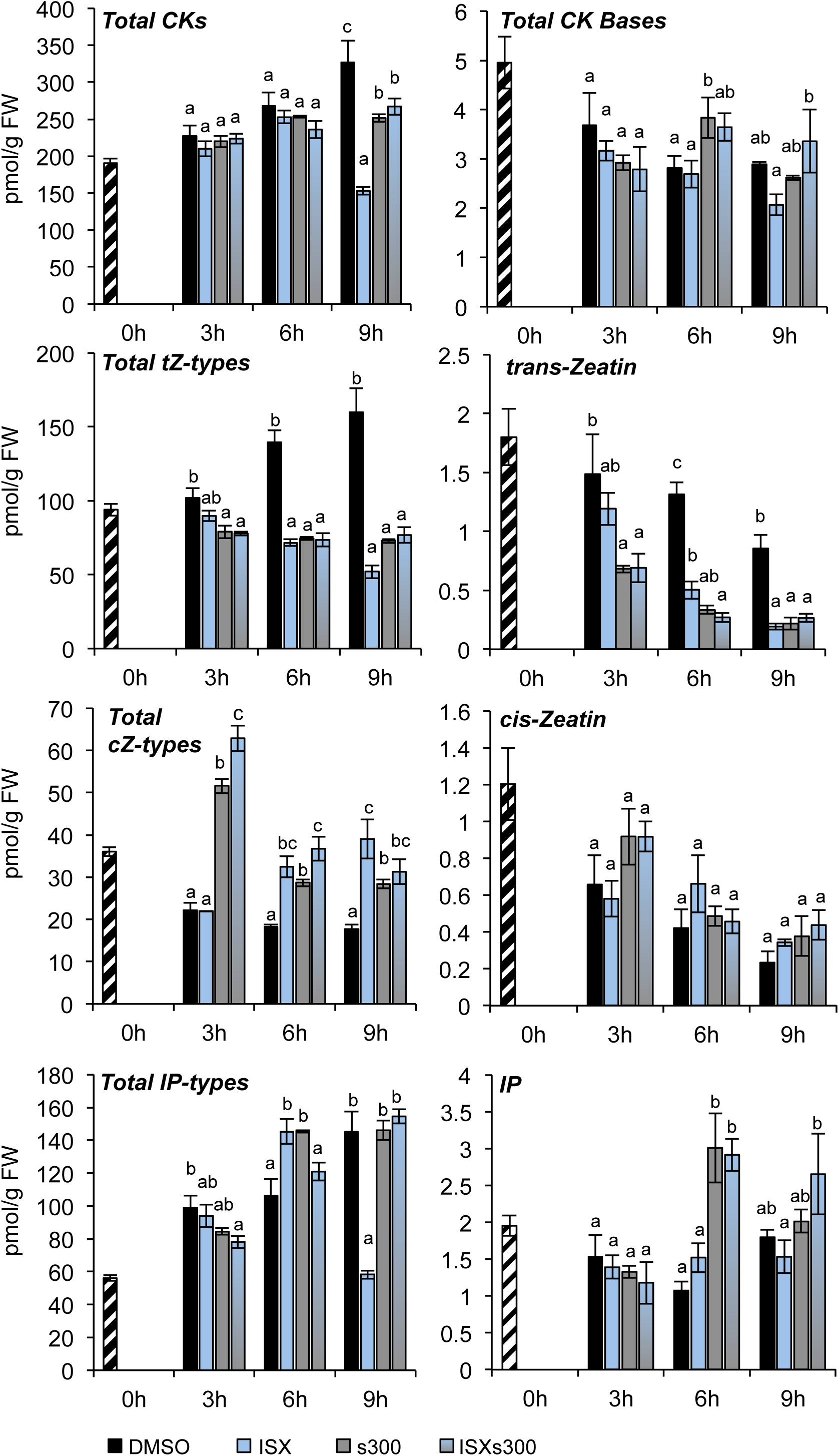
Cytokinin measurements detect differential effects of ISX and osmoticum treatments. Col-0 seedlings were treated with DMSO (mock), Isoxaben (ISX), Sorbitol (s300) or ISXs300. Total amounts of cytokinin types and corresponding bases are shown. Each column represents data deriving from 2 independent experiments with 3 biological repeats each. Error bars are based on standard deviation. One-way ANOVA and Tukey’s HSD test (α = 0.05) were used to analyze differences within single time points and different letters denote significant differences.

The measurements showed that ISX-treatments cause reductions in the levels of several cytokinins (exception *c*Z types), while s300 treatments had opposite effects (including attenuation of ISX-treatment effects). Interestingly the changes in *i*P-type levels in the ISX and /or s300 treated seedlings seem to be particularly similar to changes observed in *CYCD3;1* transcript levels.

### CKX2 and CKX3 influence osmo-sensitive effects of ISX treatments on cell cycle regulation

The results from gene expression analysis, *t*Z treatments and CK profiling suggested that ISX- and osmoticum-induced effects on *CYCD3;1* and *CYCB1;1* transcript levels may be mediated through a cytokinin-dependent mechanism. We therefore investigated if cytokinin-based signaling is regulating the ISX- and/or osmoticum-induced effects by quantifying *CYCD3;1* and *CYCB1;1* transcript levels after 9h in single, double and multiple KO seedlings for cytokinin perception and signaling elements including: *ARABIDOPSIS HISTIDINE KINASE 1 (AHK1), AHK2, AHK3, AHK4/CRE1*, *HISTIDINE-CONTAINING PHOSPHOTRANSFER PROTEIN 1 (AHP1), AHP2, AHP3, AHP4, AHP5* and *ARABIDOPSIS RESPONSE REGULATOR 1 (ARR1), ARR10, ARR12* (Zürcher & Müller, 2016). *CYCB1;1* and *CYCD3;1* transcript levels were reduced in mock-treated *ahk2/3*, *ahk3/4*, *ahp1/2/3/4/5* and *arr1/10/12* seedlings (Sup. Fig. S4a-d). This suggests that manipulation of cytokinin signaling elements affects basal expression of cell cycle genes. ISX-treated *ahk1*, *ahk2/3*, *ahk3/4*, *ahk4*, *ahp1/2/3/4/5* and *arr1/10/12* seedlings showed similarly reduced transcript levels as observed in Col-0. *CYCB1;1* levels in s300-and ISX/s300-treated *ahk2/3* seedlings seem to differ from Col-0. To address possibly existing redundancy between AHKs and avoid consequences of growth defects in *AHK*-triple KO plants we repeated some of the experiments with added co-treatments of LGR991, an AHK4 inhibitor/AHK3 competitor (Nisler *et al*., 2010). Col-0, *ahk2/3* or *ahk3/4* seedlings were treated with LGR-991 and the same experiments as before were performed (Sup. Fig. 4c). In LGR991/ISX-treated Col-0, *ahk2/3* and *ahk3/4* seedlings *CYCB1;1* and *CYCD3;1* transcript levels were still reduced. LGR991/s300-treated Col-0 seedlings exhibited lower *CYCB1;1* transcript levels than s300-treated Col-0. Both LGR-991-co-treated *ahk2/3* and *ahk3/4* seedlings still responded to the treatments but *CYCB1;1* transcript level changes were less pronounced in *ahk2/3*. These effects were not detectable for *CYCD3;1*.

Since certain cytokinin levels were changed in ISX-treated seedlings, we investigated if active degradation is responsible. Previous work suggested that KOs in individual cytokinin oxidases (CKXs) have no phenotypic effects due to redundancy (Bartrina *et al*., 2011). We used overexpression lines for *CYTOKININ OXIDASE 2* (*oeCKX2*, localized in the apoplast) and *3* (*oeCKX3*, residing in the tonoplast) because they also enabled us to investigate if spatial requirements may exist (Zürcher & Müller, 2016). We analyzed *CYCB1;1* and *CYCD3;1* transcript levels in *oe*CKX seedlings treated as before for 9 h. *CYCB1;1* transcript levels were similar in Col-0 and *oe*CKX seedlings upon mock- and s300-treatments (Fig. 6a). In ISX-treated Col-0 transcript levels were reduced while reductions in *oe*CKX seedlings were less pronounced. In ISXs300-treated Col-0 and *oeCKX3* seedlings transcript levels were similar to mock, while in *oeCKX2* they were similar to those observed in ISX-treated ones. *CYCD3;1* transcript levels were reduced in mock-treated *oeCKX2* seedlings. ISX-treated Col-0 seedlings exhibited reductions in *CYCD3;1* transcript levels, which were absent in *oeCKX2* (since basal expression was already affected) and less pronounced in *oeCKX3* seedlings. In s300- or ISX/s300-treated Col-0 seedlings *CYCD3;1* levels were enhanced. In similarly treated *oeCKX2* and *oeCKX3* seedlings the effects of s300 and ISX/s300 were less pronounced.

**Figure 6.**
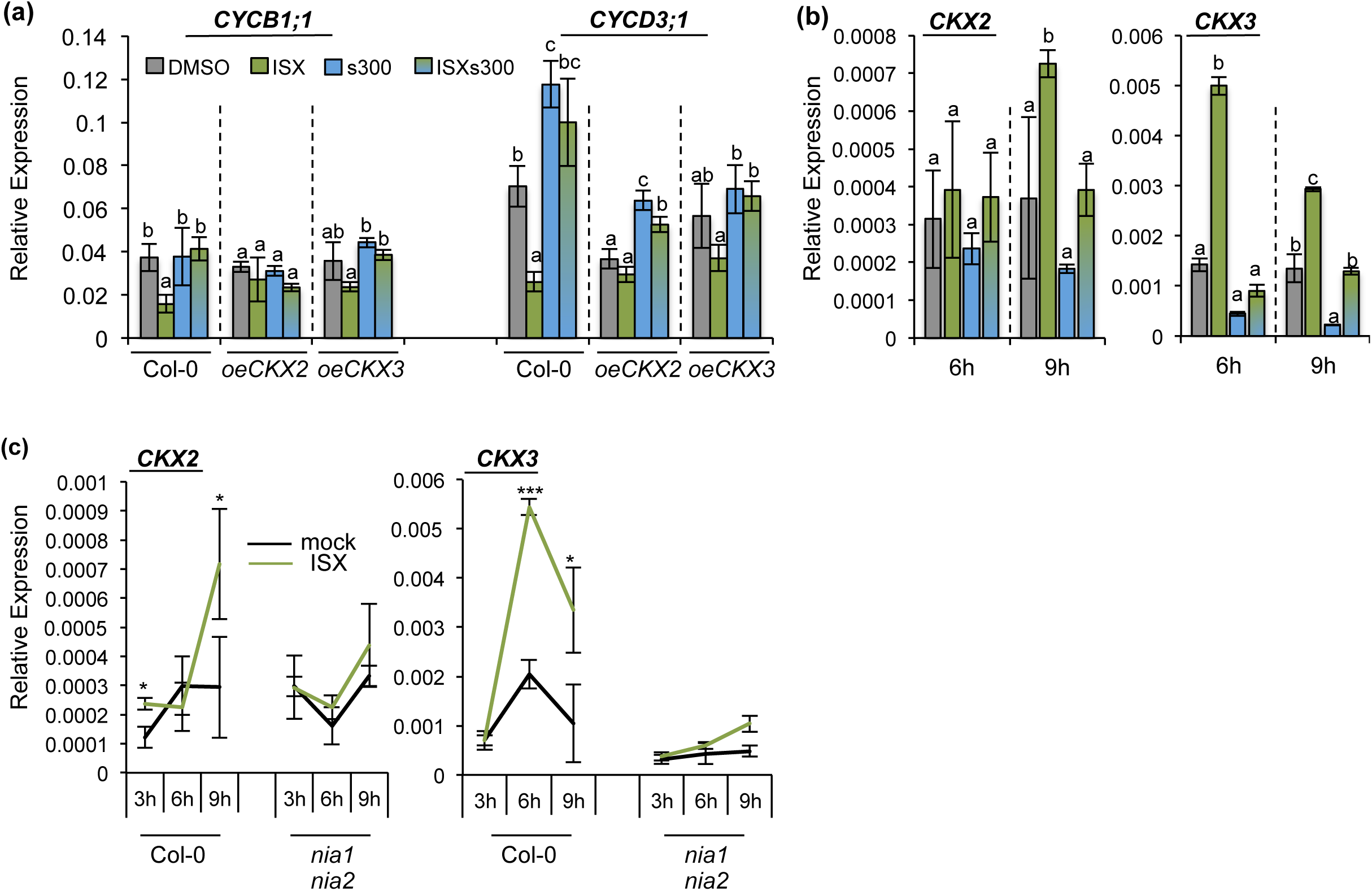
*CKX2* overexpression affects ISX- and sorbitol-induced changes in *CYCD3;1* transcript levels. **a)** Col-0, *CKX2* and *CKX3* overexpressing seedlings (*oeCKX2*, *oeCKX3*) were treated for 9 h with DMSO (mock), Isoxaben (ISX), Sorbitol (s300) or ISXs300. Relative *CYB1;1* and *CYCD3;1* expression was quantified by qRT-PCR. **b**) Relative *CKX2* and *CKX3* expression was quantified in Col-0 seedlings treated as in (a) for 6 or 9 h. All values are means and error bars are based on standard deviation (n=3). One-way ANOVA and Tukey’s HSD test (α = 0.05) were used to analyze the differences between treatments of each genotype (a) or time point (b). Different letters denote significant differences. **c**) Six-day old Col-0 and *nia1 nia2* seedlings were treated as in (a). Relative *CKX2* and *CKX3* expression was quantified in b). Values are means and error bars represent standard deviation (n=3). Pairwise comparison of ISX- and mock- treated samples was performed with Student’s t-test. *P < 0.05, *** P <0.001.

Next we investigated *CKX2* and *CKX3* transcript levels in Col-0 seedlings treated as before for 6 h or 9 h. *CKX2* transcript levels were significantly induced in seedlings after 9 h of ISX treatment (Fig. 6b). ISXs300-treated seedlings exhibited *CKX2* transcript levels similar to mock-treated controls. *CKX3* transcript levels were strongly increased after 6 h of ISX treatment, significantly reduced in s300-treated seedlings and similar to mock-controls in ISXs300-treated seedlings. To investigate if ISX-induced changes in *CKX2* and *CKX3* transcript levels are dependent on *NIA1/2* we performed time course experiments and investigated *CKX2* and *CKX3* transcript levels in mock or ISX-treated Col-0 and *nia1/2* seedlings. While *CKX2* and *CKX3* transcript levels increased over time in ISX-treated Col-0 seedlings the increase was not detectable in ISX-treated *nia1/2* (Fig. 6c).

Genetic manipulation of CK perception and signal translation did not affect the ISX-induced changes in *CYCB1;1* and *CYCD3;1* transcript levels, while osmoticum-induced changes in *CYCB1;1* transcript levels seem to be affected. ISX-induced expression of *CKX2* and *CKX3* was osmosensitive and dependent on *NIA1/2* activity. Manipulating *CKX2* expression affected *CYCB1;1* and *CYCD3;1* basal transcript levels and responses to ISX and s300 treatments. These results suggest that ISX- and osmoticum treatments modify *CKX2* and *CKX3* expression, which in turn seems to affect preferentially *CYCD3;1* transcript levels.

## Discussion

Plant cell wall metabolism and cell cycle activity are essential components of development, immunity and abiotic stress (Zhao & Dixon, 2014; Polyn *et al*., 2015; Zhao *et al*., 2017; Bacete *et al*., 2018). Here we determined how CWD (caused by cellulose biosynthesis inhibition) affects cell cycle progression and investigated the regulatory mechanism.

Phenotypic analysis of seedling root growth indicated that ISX-induced growth inhibition is attenuated by osmoticum. Analysis of RAM length and cell number showed that the osmoticum effect is restricted to attenuation of RAM length (cell size), while RAM cell number (cell division) was still arrested in ISX/s300-treated RAMs. Since cellulose is an essential element of the cell walls separating both daughter cells after mitosis, complete inhibition of cell division is to be expected and similar to effects observed in mutants affecting cellulose production (Nicol & Hofte, 1998; Miart *et al*., 2014; Chen *et al*., 2018). These results indicated that two different processes respond to ISX-treatment, osmo-sensitive cell elongation and non-sensitive cell division.

Analysis of ISX-treated Arabidopsis seedlings expressing the *pCYCB1;1::CYCB1;1 D-box-GUS* reporter suggested that after 9 h *CYCB1;1* transcript levels are reduced by ISX and attenuated by osmotic support both in shoot and root apical meristems. qRT-PCR-based expression analysis showed that *CYCB1;1* and *CYCD3;1* transcript levels were reduced by ISX treatment and similar to mock-levels if osmoticum was added, while *CDKB2;1* transcript levels did not change, suggesting *CDKB2;1* expression is not responsive to the treatments. The changes in transcript levels are apparently not induced by cell division inhibition (since cell division is still inhibited in ISX/s300 RAMs) but occur in response to a stimulus generated by ISX and sensitive to osmoticum treatment. In yeast and other plant species similar effects of osmoticum on cell cycle and survival upon exposure to CWD have been reported (Iraki *et al*., 1989; Levin & Bartlett-Heubusch, 1992).

We performed a genetic analysis and found that KOs in genes implicated in CWI maintenance do not affect the ISX-induced changes to *CYCD3;1* and *CYCB1;1* transcript levels. Previous work suggested that THE1 might modulate cell elongation in response to ISX and during organ expansion (Hematy *et al*., 2007; Fujikura *et al*., 2014). This suggests that the KOs tested here affect genes, which are also required to control ISX-effects on cell elongation but not on cell division. It also suggests the existence of a second pathway, regulating the effects on *CYCD3;1* and *CYCB1;1* transcript levels. However, both pathways respond to the same osmosensitive stimulus as indicated by the similar effects of osmoticum on CWD-induced JA, SA, lignin and *CYCD3;1* and *CYCB1;1* transcript levels (Denness *et al*., 2011).

As part of the genetic analysis we also examined *nia1/2* seedlings. Differences in the transcriptome level between Col-0 and *nia1/2* seedlings suggested that fundamental metabolic differences exist, in accordance with NIA1/2 involvement in nitrogen metabolism (Chamizo-Ampudia *et al*., 2017). However, *CYCB1;1* and *CYCD3;1* transcript levels and seedling root lengths were similar in mock-treated Col-0 and *nia1/2* seedlings, suggesting that global metabolic differences have a limited impact on the processes examined here. In these seedlings neither ISX-dependent reduction of *CYCB1;1* and *CYCD3;1* transcript levels nor induction of JA/SA/lignin accumulation were detectable. Simultaneously NIA1/2 are required to mediate the THE1-dependent responses (based on the *the1-4 nia1/2* results), implying that NIA1/2-dependent processes are involved in both THE1-dependent and -independent processes.

*nia1/2* seedlings have been used as genetic tool to study NO-based signaling processes and been identified as important regulators of abiotic stress responses (Xie *et al*., 2013; Vicente *et al*., 2017). NO has also been suggested to regulate and be regulated by cytokinins suggesting the existence of a regulatory feedback loop between the two (Tun *et al*., 2008; Feng *et al*., 2013; Liu *et al*., 2013; Shen *et al*., 2013). In Col-0 seedlings, treated jointly with increasing concentrations of *t*Z and ISX, ISX-induced reductions in *CYCD3;1* transcript levels were alleviated in a concentration dependent manner by *t*Z. In *nia1/2* seedlings, ISX-induced reductions in transcript levels were absent while *tZ* treatment effects were still detectable, implying that the effects do not require functional NIA1/2. These results suggested that ISX-effects on *CYCD3;1* transcript levels and activity might be mediated through changes in cytokinin levels, similar to previously reported regulation of the gene by cytokinins (Scofield *et al*., 2013).

Measurements of cytokinin levels in ISX-, osmoticum or ISX/osmoticum-treated Col-0 seedlings showed that individual treatments had opposite effects on certain cytokinin amounts while the combined treatments resulted in amounts similar to mock controls. These results were similar to what we observed primarily for *CYCD3;1* transcript levels. Intriguingly *i*P type level changes were particularly similar to *CYCD3;1* transcript level changes, suggesting they may be responsible for the effects observed. Experiments with maize exposed to osmotic stress have also shown that osmotic stress affects cytokinin levels, but the experimental conditions differed, making comparisons difficult (Vyroubalová *et al*., 2009). We performed a genetic analysis to determine if cytokinin signaling is required for the effects of ISX and osmoticum treatments on *CYCD3;1* and *CYCB1;1* transcript levels. Mock-treated *ahk2/3*, *ahk3/4*, *arr1/10/12* seedlings had lower basal *CYCD3;1* transcript levels indicative of cytokinin influence on *CYCD3;1* expression and cell cycle activity. *ahk2/3* seedlings also exhibited changes in transcript levels upon treatment with osmoticum only, with the underlying reason remaining to be resolved. However, in all the genotypes examined (also in combination with LGR-991 treatments), ISX-induced reductions in transcript levels were still detectable, suggesting that intact cytokinin signaling is not required. However, expression of two genes encoding cytokinin-degrading enzymes (*CKX*2, *CKX3*) was induced by ISX treatment in a *NIA1/2*-dependent and osmo-sensitive manner.

Characterization of *CYCD3;1* and *CYCB1;1* transcript levels in ISX-and / or osmoticum-treated *oeCKX2* and *3* seedlings suggested that *CKX2* and *CKX3* overexpression affects ISX- and osmoticum-induced changes. Bearing in mind that CKX enzymes cleave the isoprenoid side chain of *N*^6^-(Δ^2^-isopentenyl) adenine (iP), *trans*- and *cis*-zeatin (*t*Z, *c*Z) and their ribosides (iPR, *t*ZR and *c*ZR), leading to adenine/adenosine and the corresponding aldehyde, both CKX2 and 3 could be responsible for the changes in cytokinin levels observed (Kopečný *et al*., 2016). The results suggest that ISX-induced CWD causes increased expression of *CKX2* and *3* through NIA1/2-dependent processes. Increased expression of *CKX2* and *CKX3* could lead to reductions in certain cytokinins, which in turn reduces transcript levels of *CYCD3;1* and cell cycle progression.

We have integrated the results presented here into a model describing the mechanism regulating adaptation of cell elongation and cell cycle activity to CWD (Figure 7). CWD is perceived by an unknown, osmo-sensitive CWI sensor, which activates at least two different signaling cascades: one *THE1*-dependent (required for induction of JA/SA/lignin production and possibly inhibition of cell elongation) and a second, *THE1*-independent one, implying that *THE1* does not encode a unique CWI sensor. The *THE1*-independent one would be responsible for the effects on *CYCD3;1* and *CYCB1;1* transcript levels. *NIA1/2-*dependent processes are required to mediate both *THE1*-dependent (JA/SA/lignin) and -independent responses. *NIA1/2* seem also responsible for controlling *CKX2* and *CKX3* expression. Changes in *CKX2* and *3* transcript levels are correlated with reductions or increases in certain cytokinin levels upon ISX, osmoticum and co-treatments of both. This correlation suggests that the changes, achieved by controlled degradation through CKX2 and CKX3, could be responsible for modifying *CYCD3;1* expression, which in turn might adapt cell cycle progression to the state of the cell wall (CWI). While cytokinin signaling redundancy cannot be excluded (based on our data), in this model intact cytokinin perception and signaling are actually not required. The reason being that regulation of *CKX2* and *3* by *NIA1/2*-mediated processes would cause reductions in cytokinin levels. These would in turn lead to reduced cell cycle activity, thus bypassing cytokinin signaling and explaining our results with cytokinin signaling mutants.

**Figure 7.**
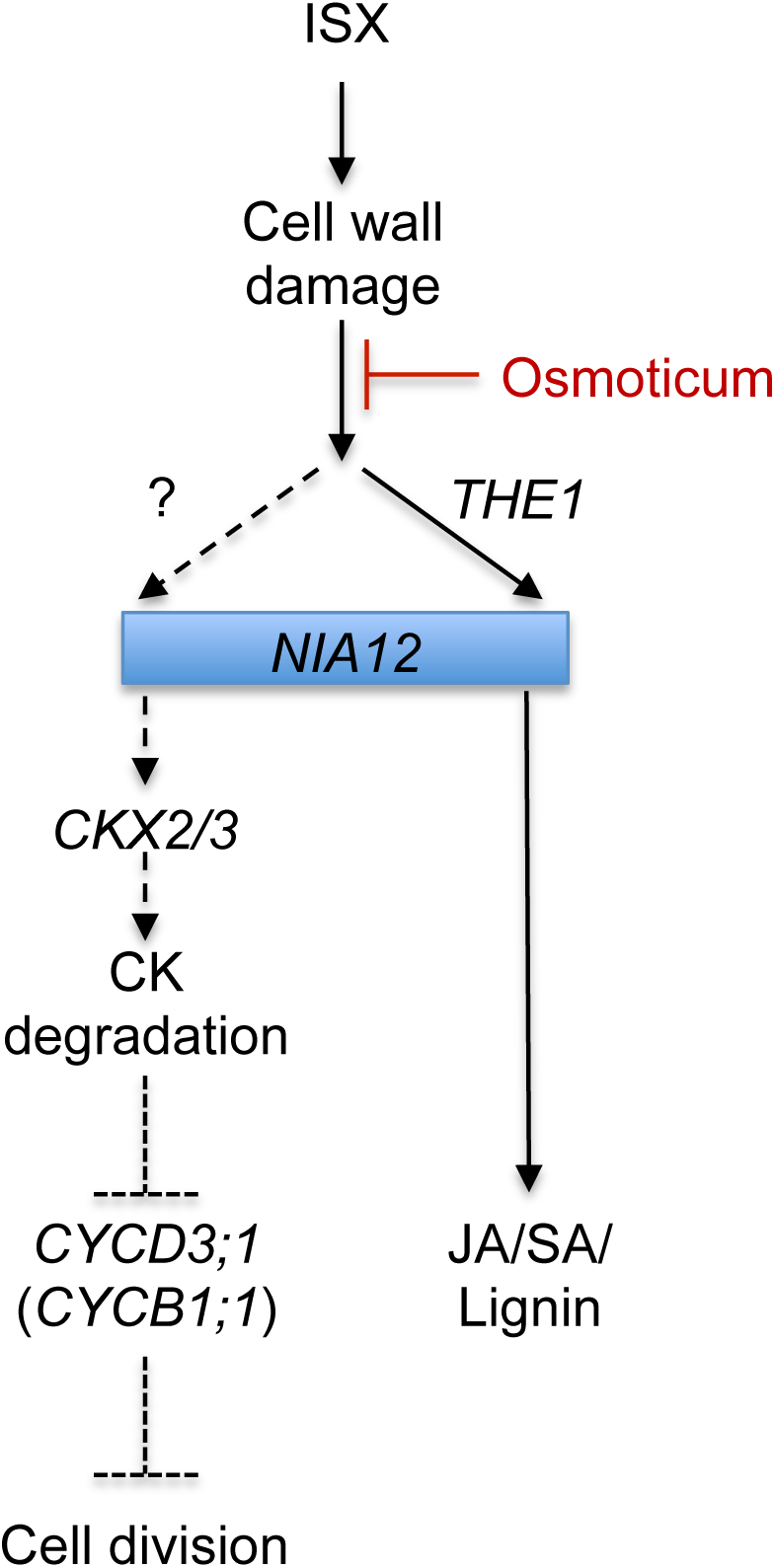
The mechanism coordinating CWI with cell cycle progression and other ISX-induced responses. ISX-induced CWD is osmo-sensitive. CWD is perceived by an osmo-sensitive mechanism, which then activates two signaling pathways. One is *THE1*-dependent (required for induction of JA/SA/lignin accumulation). A second *THE1*-independent one, which is responsible for the changes observed in expression of *CYCD3;1* (and *CYCB1;1*). *NIA1 NIA2-* dependent processes are required to mediate both *THE1*-dependent and independent responses. NIA1 NIA2 seem also responsible for controlling expression of *CKX2* and *CKX3* upon ISX treatment. Expression of these *CKX* genes is enhanced by ISX and reduced by osmotic treatments, which might correlate with reductions in certain cytokinins (CKs) upon ISX- and increases upon treatments with osmoticum. Changes in certain CKs, could be responsible for modifying expression of *CYCD3;1*, which in turn controls cell cycle progression. Dashed lines indicate possible regulatory relationships.

In the experiments performed, osmoticum reduced ISX-induced effects (cell elongation, JA/SA/lignin accumulation, transcript and cytokinin levels). Such effects of turgor on cell cycle activity have been observed earlier in plants, yeast and been discussed as important regulators of cell cycle progression (Iraki *et al*., 1989; Levin & Bartlett-Heubusch, 1992; Sablowski, 2016). Conceivable explanations for the osmoticum effect are that CWD involves plasmamembrane displacement vs. the cell wall, changes in surface tension of the cell wall itself or simply changes in the optimal state of the plasmamembrane (stretching/shrinking), which could be detected by dedicated sensor proteins similar to the mechanisms in place in yeast or animal cells (Heinisch *et al*., 2010; Austen *et al*., 2015; Hamant & Haswell, 2017). Elucidation of these sensor systems will provide the mechanistic insights necessary to understand how the CWI is monitored in plants and downstream responses activated.

## Acknowledgements

The authors thank Hana Martínková and Ivan Petřík for their help with cytokinin analyses and John Mansfield for comments on the manuscript.

Financial and/or technical support from HORIZON2020 (SugarOsmoSignaling) (T.E.), the EEA grant 7F14155 CYTOWALL (N.G.B./T.H./J.H) and Ministry of Education, Youth and Sports of the Czech Republic (National Program for Sustainability I, grant no. LO1204), are gratefully acknowledged.

## Supplementary Information

**Table S1:** Overview of genotypes employed

**Table S2:** Overview of PCR-Primers used

**Table S3:** GO Analysis of candidate genes identified in transcriptomics experiment part a

**Table S4:** GO Analysis of candidate genes identified in transcriptomics experiment part b

**Table S5:** GO Analysis of candidate genes identified in transcriptomics experiment part c

**Table S6:** Overview of candidate genes exhibiting statistically significant differences between treatments and genotypes.

**Figure S1:**
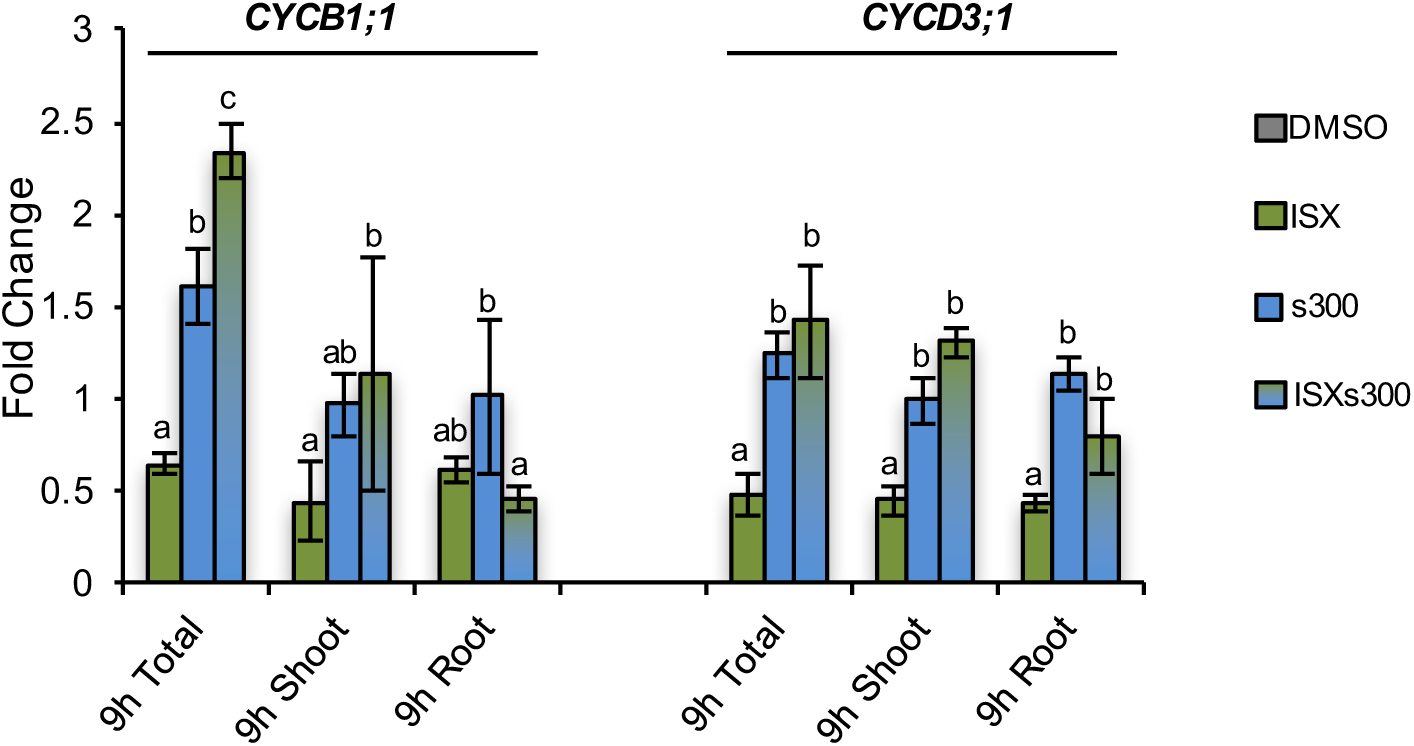
*CYCB1;1* and *CYCD3;1* expression levels are similar in roots and shoots. To separate between shoots and roots, Col-0 seeds were placed on sterile pads fixed inside sterile bottles containing medium. After six days the medium contained in the flasks was replaced supplemented with DMSO, Isoxaben (ISX), Sorbitol (s300) or a combination of both (ISXs300) for 9 h. Relative expression of *CYCB1;1* and *CYCD3;1* was determined in whole seedlings (total), shoots or roots by qRT-PCR. Data are shown relative to mock (DMSO) treatment (black dashed line). Values are means and error bars are based on standard deviation (n=3). Different letters denote statistically significant differences according to one-way ANOVA and Tukey’s HSD test (α = 0.05).

**Figure S2.**
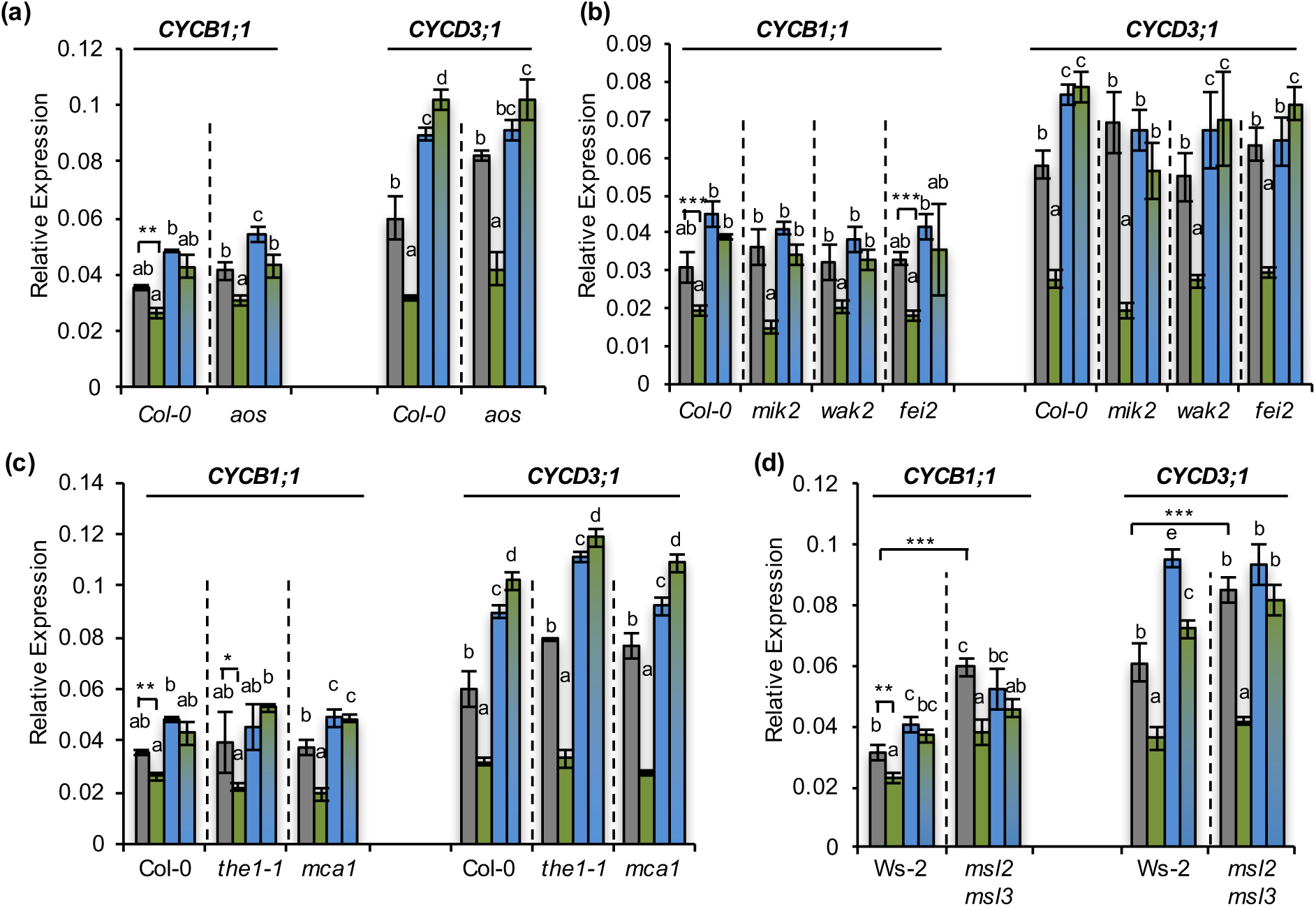
Effects of ISX-treatments on *CYCB1;1* and *CYCD3;1* transcript levels in CWI maintenance KOs. Relative expression of *CYCB1;1* and *CYCD3;1* was determined in different genotypes after mock-(DMSO), Isoxaben-(ISX), Sorbitol-(s300) or ISXs300 treatment for 9 h. a) Col-0, *aos*; **b)** Col-0, *mik2*, *fei2*, *wak2*; c) Col-0, *the1-1*, *mca1*; d) Ws-2, *msl2 msl3.* Values in (a-d) are means and error bars are based on standard deviation (n=3 independent experiments). One-way ANOVA and Tukey’s HSD test (α = 0.05) were used to analyze the differences between treatments of each genotype and different letters denote significant differences. Pairwise comparisons in (a-d) have been performed as indicated with Student’s t-test. * P < 0.05, ** P <0.01, *** P <0.001.

**Figure S3.**
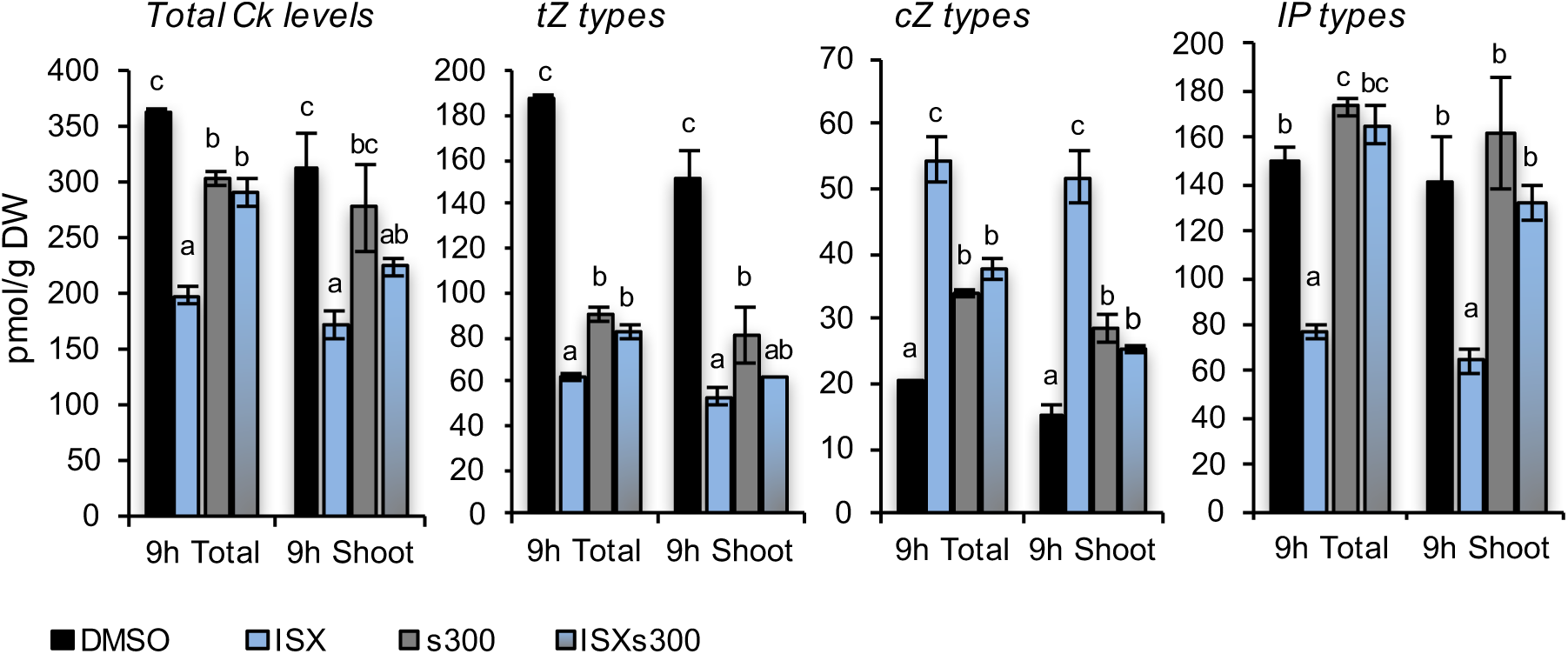
ISX and osmoticum treatments affect cytokinin levels in whole seedlings and aerial parts similarly. Total Cytokinin levels and Cytokinin types were analyzed in Col-0 seedlings treated with DMSO, isoxaben (ISX), Sorbitol (s300, 300mM) or a combination thereof (ISXs300) for 9 h. Values are means and error bars are based on standard deviation (n=4). Different letters denote significant differences between treatments of whole seedlings or shoots only, according to one-way ANOVA and Tukey’s HSD test (α = 0.05).

**Figure S4.**
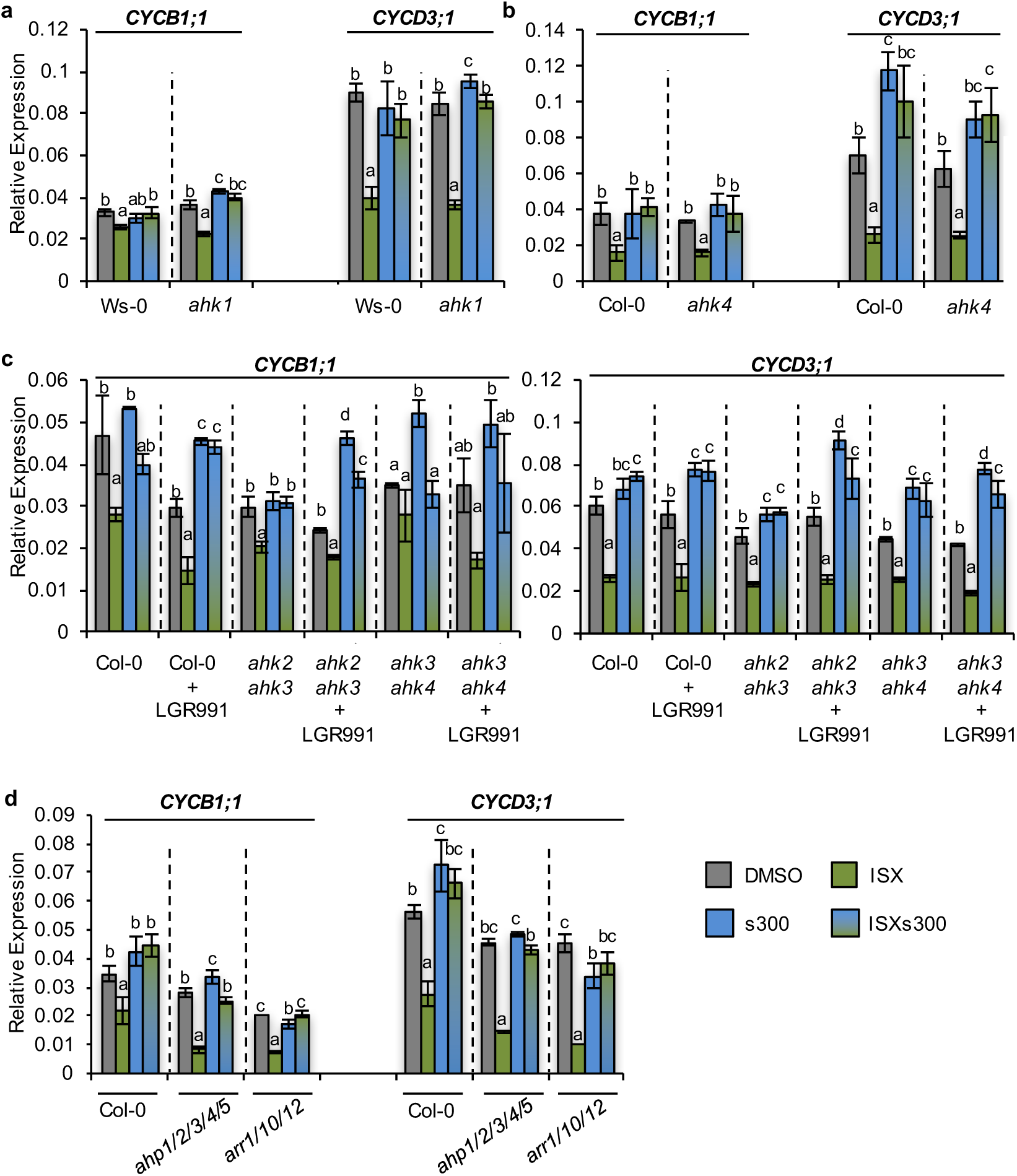
*CYCB1;1* and *CYCD3;1* transcript levels in seedlings deficient in cytokinin signaling. Seedlings of different genotypes were treated for 9h with DMSO, Isoxaben (ISX), Sorbitol (s300) or a combination thereof (ISXs300) for 9h. **a**) Ws-0, *ahk1;* **b**) Col-0, *ahk4;* **c**) Col-0, *ahk2 ahk3* and *ahk3 ahk4* seedlings were treated with DMSO, Isoxaben (ISX), Sorbitol (s300) or both (ISXs300) +/- 20 µM LGR-991. **d**) Col-0, *ahp1/2/3/4/5* and *arr1/10/12* seedlings were treated as in a. Error bars are based on standard deviation (n=3). Letters are shown according to one way ANOVA. Tukey’s HSD test (α = 0.05).

